# Multiple ammonium transporters in fission yeast are coordinated by transcriptional and localization regulation in response to nitrogen starvation

**DOI:** 10.1101/2025.10.25.684503

**Authors:** Yukiko Nakase, Ng Shet Lee, Ayu Shibatani, Takeru Yomogita, Yuichi Morozumi, Daisuke Watanabe, Hiroshi Takagi, Kazuhiro Shiozaki

## Abstract

Ammonium is a preferred nitrogen source for microorganisms. We investigated the regulation of ammonium transporters Amt1, Amt2, and Amt3 in the fission yeast *Schizosaccharomyces pombe*. Expression of the *amt1*^+^ gene increases under nitrogen starvation as well as in the TORC1-deficient mutant *tor2-287*, suggesting the negative regulation of *amt1*^+^ by TORC1. This regulation depends on the GATA transcription factor Gaf1, whose phosphorylation is regulated by Ppe1 and PP2A phosphatases. Ppe1 is required for the nuclear accumulation of Gaf1, partly contributing to the controlled expression of *amt1*^+^. On the other hand, PP2A phosphatase is essential for the *amt1*^+^ induction upon starvation, indicating distinct roles for Ppe1 and PP2A in Gaf1 regulation. Nitrogen starvation also promotes the plasma membrane translocation of Amt1 and Amt2 in a manner dependent on the Tsc-Rhb1 pathway. Intriguingly, the *amt1Δ amt2Δ tscΔ* mutant is more sensitive to low-ammonium conditions than the *amt1Δ amt2Δ amt3Δ* triple mutant. Thus, there may be an unidentified ammonium transporter whose plasma membrane localization is also dependent on the Tsc-Rhb1 pathway.

## Introduction

The Target of Rapamycin (TOR), a serine/threonine protein kinase highly conserved among eukaryotes, is a pivotal regulator of cellular growth in response to extracellular nutrients (Gonzalez and Rallis, 2017; Valvezan and Manning, 2019). Biochemical analysis identified two types of cellular protein complexes containing the TOR kinase, TOR complex 1 (TORC1) and TOR complex 2 (TORC2). TORC1 and TORC2 are responsive to distinct extracellular and intracellular inputs and control different sets of cellular processes (Loewith et al., 2002; Wullschleger et al., 2006). TORC1 promotes anabolic processes essential for proliferation, such as protein translation and ribosome biogenesis, while inhibiting autophagy (Schmelzle and Hall, 2000). Although only one TOR kinase is present in most eukaryotic species, two TOR kinases, Tor1 and Tor2, exist in the well-studied yeast species *Saccharomyces cerevisiae* and *Schizosaccharomyces pombe* (Heitman et al., 1991; Otsubo and Yamamato, 2008; Weisman and Choder, 2001). In the fission yeast *S. pombe*, TORC1 comprises the catalytic subunit Tor2 and its regulatory subunits, including Mip1 (a human Raptor homolog) and Wat1 (a human mLst8 homolog). TORC1 is regulated through a signaling network that contains Rhb1, a Rheb-family GTPase, and the heterodimeric Gtr1-Gtr2 GTPases orthologous to human RagA/B-RagC/D; in response to changes in nutrient availability, the TORC1 activity is controlled by the counteracting actions of the positive regulator Rhb1 and the negative regulator Gtr1-Gtr2 heterodimer (Chia et al., 2017; Mach et al., 2000; Urano et al., 2005; Uritani et al., 2006). Rhb1 is negatively regulated by the Tsc1-Tsc2 complex, which acts as a GTPase-activating protein (GAP) (Matsumoto et al., 2002; Nakase et al., 2006). In humans, the *TSC1* and *TSC2* genes have been identified as causative for tuberous sclerosis, and their orthologues are conserved in fission yeast but not in budding yeast. In addition to TORC1, the fission yeast Tsc complex-Rhb1 pathway controls the translocation of amino acid transporters to the plasma membrane in response to nitrogen starvation (Aspuria and Tamanoi, 2008; Nakase et al., 2013). In the downstream of the Tsc-Rhb1 pathway, the E3 ubiquitin ligase Pub1 and the β-arrestin-like protein Any1 form a complex that regulates the localization of amino acid transporters via ubiquitination, thereby controlling the cellular amino acid uptake.

Ammonium, a vital nitrogen source for many microorganisms, induces the robust activation of fission yeast TORC1 and the phosphorylation of its substrates, including Psk1, a homolog of mammalian S6K1 (Nakashima et al., 2012). In response to nitrogen starvation, TORC1 is inactivated, resulting in the dephosphorylation of Psk1. Feeding ammonium to the starved cells swiftly brings about the reactivation of TORC1 and the Psk1 phosphorylation. Thus, ammonium is a potent inducer of TORC1 signaling in fission yeast, though the molecular mechanism of how ammonium triggers TORC1 activation remains unknown.

The integral membrane proteins of the Mep-Amt-Rh family mediate ammonium transport across the plasma membrane in diverse cell types, ranging from bacteria to mammals (Biver et al., 2008; Huang and Peng, 2005; Marini et al., 1997). Among the family members are Mep1, Mep2, and Mep3 in budding yeast, of which transporter activity is regulated by TORC1 signaling. Under nitrogen-replete conditions, TORC1 phosphorylates and inhibits the Npr1 kinase. In response to nitrogen starvation and resultant TORC1 inhibition, activated Npr1 phosphorylates Par32/Amu1, a direct inhibitor of Mep1 and Mep3; the phosphorylated Par32/Amu1 is released from Mep1 and Mep3, leading to the activation of these transporters (Boeckstaens et al., 2015; Varlakhanova et al., 2018). Npr1 also directly phosphorylates Mep2 and enhances its ammonium transporter activity (Boeckstaens et al., 2014).

Budding yeast TORC1 also transcriptionally controls the Mep ammonium transporters. TORC1 regulates the nuclear localization of the GATA transcription factor Gln3 (Beck and Hall, 1999; Bertram et al., 2000), which mediates the expression of the *MEP* genes (Marini et al., 1997). Upon nitrogen starvation, TORC1 is inactivated, resulting in the activation of the Sit4 and PP2A phosphatases that dephosphorylate Gln3. The dephosphorylated form of Gln3 translocates into the nucleus, promoting the transcription of the *MEP* and other permease genes required to uptake nitrogen-containing nutrients.

The GATA transcription factors in fission yeast Gaf1 and Fep1 have DNA-binding domains and zinc finger motifs resembling those of Gln3 in budding yeast. Previous studies implicated Gaf1 in regulating sexual development upon nitrogen starvation (Kim et al., 2012) and Fep1 in iron homeostasis (Pelletier et al., 2003). Like Gln3 in budding yeast, the nuclear accumulation of Gaf1 is induced upon nitrogen starvation through a mechanism mediated by TORC1 and the Ppe1 protein phosphatase, a fission yeast counterpart of Sit4 in budding yeast (Laor et al., 2015). This regulated nuclear localization of Gaf1 has been proposed to be responsible for the starvation-induced transcription of amino-acid permease genes and *isp7^+^*, a gene required for the permease expression. On the other hand, it has not been studied whether Gaf1 and/or Fep1 are involved in the regulation of the ammonium transporter genes in fission yeast *amt1*^+^, *amt2*^+^, and *amt3*^+^ (Mitsuzawa, 2006), which are orthologous to the budding yeast *MEP* genes.

As ammonium is a favorable nitrogen source and a strong promoter of the TORC1 activity in yeast, its cellular uptake is likely a crucial regulatory step in response to the nutrient. In this study, we have delved into the regulation of the Amt1, Amt2, and Amt3 ammonium transporters in fission yeast. The three Amt proteins are differentially expressed, with Amt2 being a major ammonium transporter. Among the three *amt*^+^ genes, only *amt1*^+^ is transcriptionally induced in response to nitrogen starvation and, thus, TORC1 inactivation. This regulated *amt1*^+^ expression is dependent on the GATA transcription factor Gaf1, but, unexpectedly, its Ppe1-dependent nuclear accumulation upon starvation is not essential for the induced *amt1*^+^ expression. On the other hand, the PP2A phosphatase activity is essential for the starvation-induced *amt1*^+^ expression, suggesting that the Ppe1 and PP2A phosphatases have distinct roles in the Gaf1 regulation. We have also found that the plasma membrane targeting of Amt1 and Amt2 is regulated in response to nitrogen starvation by the Pub1-Any1 complex through the Tsc-Rhb1 pathway. Interestingly, the *amt1Δ amt2Δ tscΔ* triple mutants fail to grow on low ammonium conditions, where the *amt1Δ amt2Δ amt3Δ* triple mutant can grow. This observation implies that fission yeast has one or more unknown ammonium transporters whose membrane targeting is also dependent on the Tsc-Rhb1 pathway.

## Results

### Amt proteins are essential for ammonium uptake under low ammonium conditions

Three ammonium transporters, Amt1, Amt2, and Amt3, have been reported in the fission yeast *S. pombe*; they are highly homologous to the Mep proteins in the budding yeast *S*. *cerevisiae* (Marini et al., 1997; Mitsuzawa, 2006). The sequence homology and the mutant phenotypes suggested that the Amt proteins function as ammonium transporters. We constructed a series of *amt* deletion strains and evaluated their growth in different ammonium concentrations by spotting the serial dilutions of the mutant cell cultures. None of the single deletion strains tested exhibited an apparent growth phenotype on the standard EMM medium containing 93 mM ammonium, and on that with ammonium reduced to 10 mM or 0.1 mM (Fig. 1A). On the other hand, the *amt1Δ amt2Δ* double and the *amt1Δ amt2Δ amt3Δ* triple mutants exhibited severe growth defects were under the limited ammonium conditions (10 mM and 0.1 mM).

**Figure 1.**
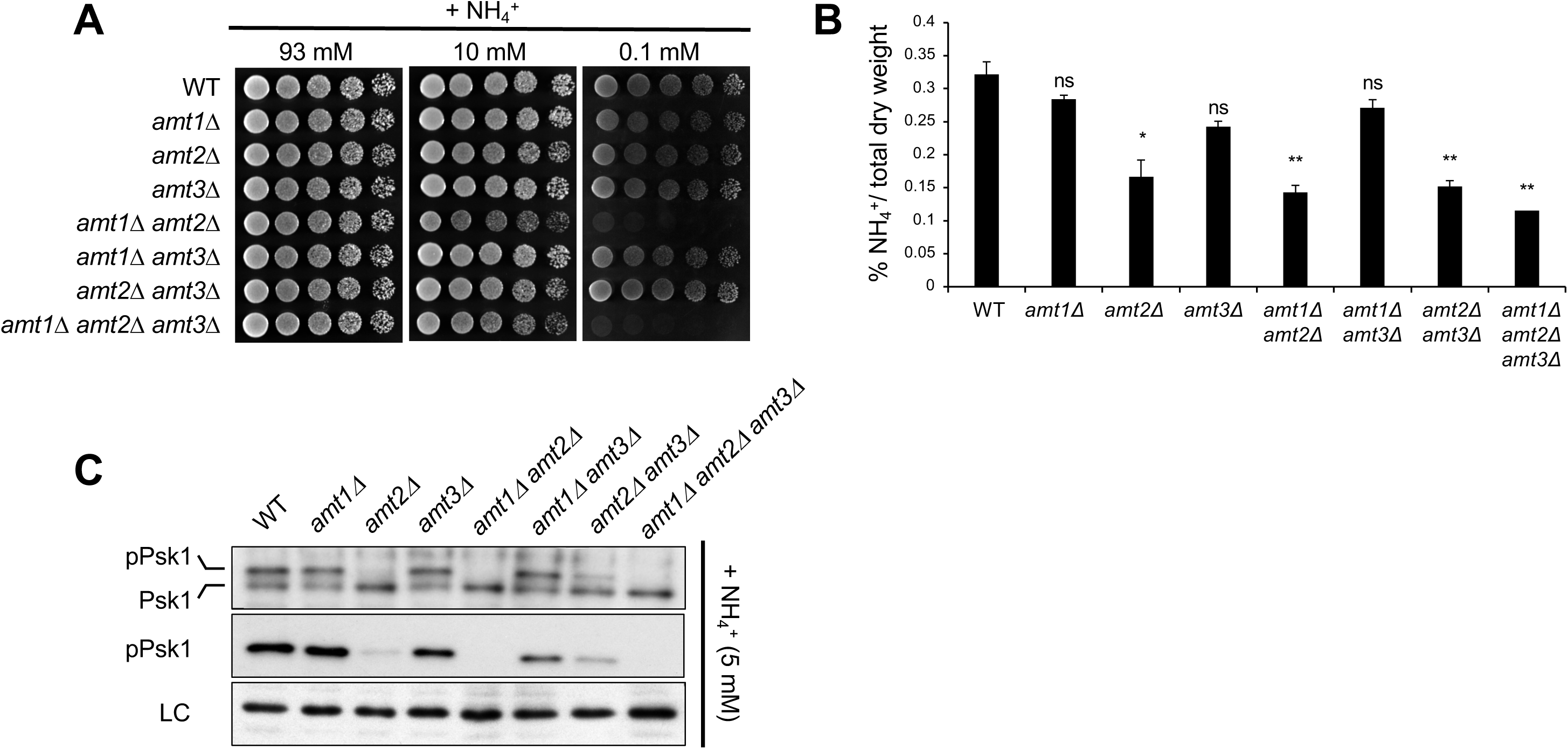
Fission yeast ammonium transporters are required for growth under low ammonium conditions. (A) Amt1 and Amt2 are important for growth under low ammonium conditions. Cells of the indicated strains were cultured in EMM medium and their serial dilutions were spotted onto EMM agar plates supplemented with different concentrations of ammonium (0.1 mM, 10 mM and 93 mM) and further incubated at 30 ℃ for 3 days. (B) Intracellular ammonium is reduced in *amt* mutants. Cells of the indicated strains were grown in EMM medium supplemented with 10 mM ammonium. The intracellular ammonium levels were analyzed by HPLC. Mean and SEM of three biological replicates are shown. Asterisks indicate t. test values; non-significant (*ns*), p < 0.05 (*\**) and p < 0.01 (*\*\**). (C) Loss of *amt* genes lead to TORC1 inhibition under low ammonium conditions. Cells of the indicated strains were grown in EMM medium supplemented with 5 mM ammonium at 30 °C. The crude cell lysates were analyzed by immunoblotting using anti-Psk1, anti-pPsk1, and anti-Spc1 (loading control) antibodies.

To confirm that the observed growth defects are due to the defective ammonium uptake, we measured the intracellular ammonium levels in the *amt* single, double, and triple mutants. The mutant strains were cultured in liquid growth medium with 10 mM ammonium, the concentration that allows even the triple *amt* mutant to grow (Fig. 1A). The harvested cells were heated to 95°C in water, and the extracted ammonium was quantified (Materials and Methods). There was no significant difference in the intracellular ammonium levels among the wild type, *amt1Δ,* and *amt3Δ* strains (Fig. 1B). In contrast, any strains lacking the *amt2^+^* gene, such as *amt2Δ, amt1Δ amt2Δ, amt2Δ amt3Δ,* and *amt1Δ amt2Δ amt3Δ*, exhibited decreased intracellular ammonium levels; the triple deletion mutants displayed the lowest level of intracellular ammonium. These results suggest that, among the three Amt proteins, Amt2 carries a major ammonium transporter activity and that Amt1 and Amt3 contribute much less to cellular ammonium uptake.

As the activation of TORC1 is dependent on nitrogen sources, such as ammonium (Nakashima et al., 2012), the TORC1 activity should be affected when cellular ammonium uptake is limited. Therefore, we assessed the TORC1 activity in the *amtΔ* mutants grown under a low ammonium condition (5 mM) by monitoring the phosphorylation of Psk1, a known substrate of TORC1 (Nakashima et al., 2012). The phosphorylation of Psk1 by TORC1 results in reduced electrophoretic mobility of Psk1 and is also detectable using antibodies against phosphorylated human S6K1. The strains lacking the *amt2^+^* gene exhibited low phosphorylation levels of Psk1 (Fig. 1C). This *amt2Δ* phenotype was exaggerated when combined with the *amt1Δ* mutation, as observed in the *amt1Δ amt2Δ* and *amt1Δ amt2Δ amt3Δ* strains. On the other hand, we detected no significant difference in the Psk1 phosphorylation between the wild-type and *amt* mutant strains in the presence of 93 mM ammonium (Fig. S1).

Together, these results strongly suggest that the Amt proteins play a crucial role in cellular ammonium uptake, particularly under low ammonium conditions, and affect the TORC1 activity. The Amt1 and Amt2 proteins have functional redundancy, with Amt2 as a major ammonium transporter. On the other hand, the contribution of Amt3 is not necessarily apparent even in the *amt1Δ amt2Δ* background.

### The expression of Amt proteins is differentially regulated

The notable difference in the contributions of the three Amt paralogs to cellular ammonium uptake might be due to their differential expression levels. To examine the expression of the Amt proteins, we inserted the FLAG epitope sequences at the 3’ end of each *amt^+^* open reading frame so that Amt1, Amt2, and Amt3 are expressed from their chromosomal loci with the C-terminal FLAG tag. Anti-FLAG immunoblotting found that the expression of Amt1 was induced in response to ammonium starvation (Fig. 2A). The smeared bands of the Amt2 protein were detected at similar levels both in the presence and absence of ammonium. On the other hand, Amt3-FLAG was undetectable under both ammonium conditions, implying that Amt3 is expressed at a much lower level than Amt1 and Amt2. The low expression level of Amt 3 seems consistent with the observations that the *amt3Δ* mutant exhibits no apparent phenotype (Fig. 1).

**Figure 2.**
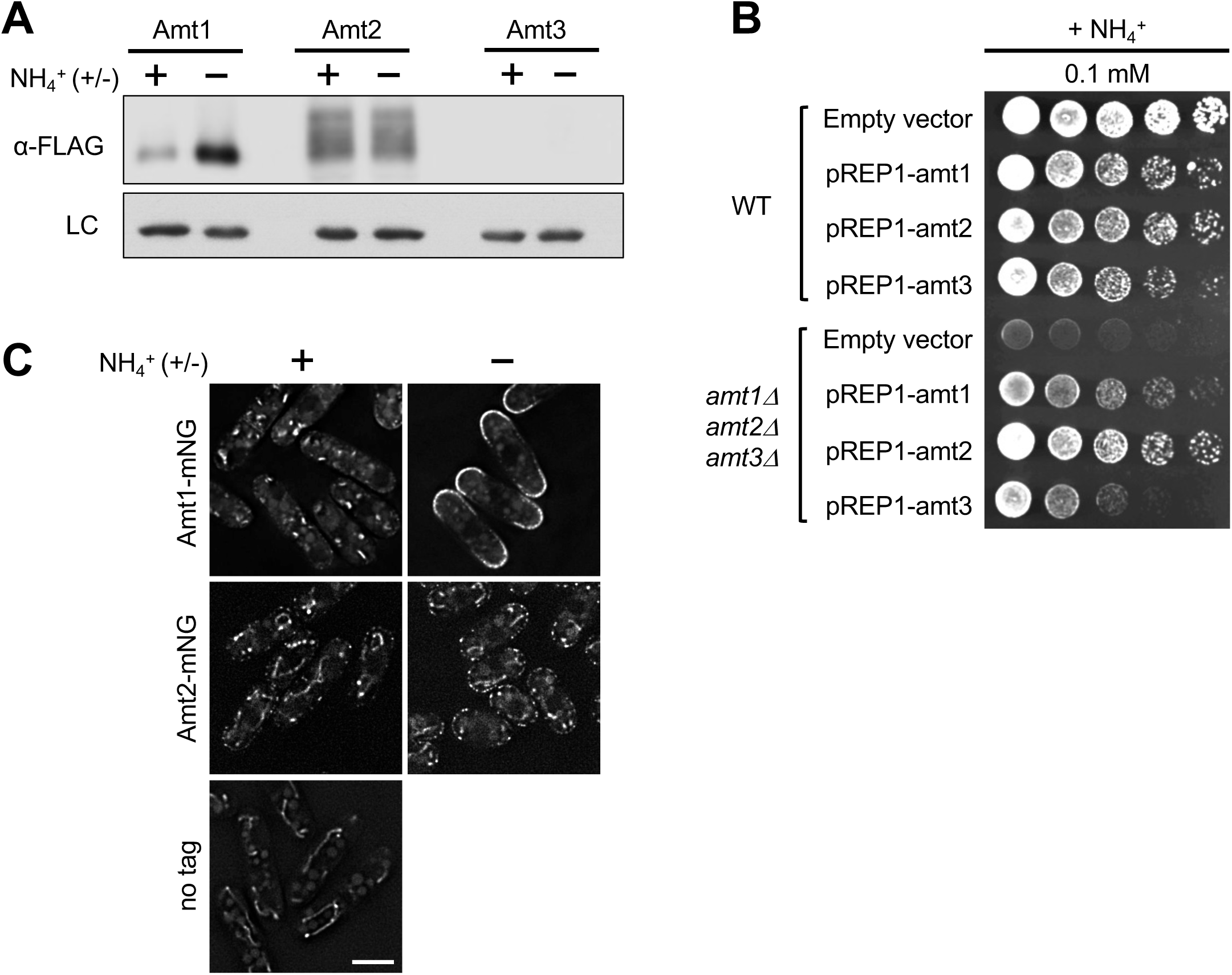
The proteins Amt1, Amt2, and Amt3 exhibit varying expression levels. (A) Amt proteins exhibit different expression levels. Cells expressing Amt1-FLAG, Amt2-FLAG or Amt3-FLAG were cultured in EMM liquid medium at 30℃ and transferred to EMM lacking ammonium for one hour. The crude cell lysates were analyzed by immunoblotting using anti-FLAG and anti-Spc1 (loading control) antibodies. (B) Overexpression of each Amt proteins complements the growth defect of triple *amt* deletion mutant. The wild-type and *amt1Δ amt2Δ amt3Δ* strains were transformed with the pREP1 plasmids carrying the *amt1^+^*, *amt2^+^* or *amt3^+^* genes, under the control of the thiamine-repressible nmt1 promoter (P*_nmt1_*). The cells were cultured in EMM liquid medium supplemented with 2 μM thiamine. To induce the overproduction of the Amt proteins, thiamine was then washed off and serial dilution of the cells were spotted on EMM agar medium with 0.1 mM ammonium and further incubated at 30 ℃ for 3 days in the absence of thiamine. (C) Amt1 localizes to plasma membrane in response to nitrogen starvation and Amt2 is constitutively present on plasma membrane. Cells expressing Amt1-mNG or Amt2-mNG were cultured in EMM at 30℃ and transferred to EMM lacking ammonium. The cells were collected at the indicated time points and observed under a fluorescence microscope. Scale bar: 5 μm.

As the loss of *amt3*^+^ exhibits no significant phenotype, it is unclear whether the gene encodes a functional ammonium transporter. We, therefore, overexpressed each Amt protein in the *amt1Δ amt2Δ amt3Δ* triple mutant under the control of the thiamine-repressible *nmt1* promoter (Maundrell, 1990). Overexpression of any one of the *amt*^+^ genes, including *amt3*^+^, complemented the growth defect of the triple *amt* deletion mutant in the presence of 0.1 mM ammonium, albeit to different degrees (Fig. 2B). Thus, not only Amt1 and Amt2 but also Amt3 appear to have the ability to function as an ammonium transporter, at least when overexpressed.

The cellular expression of the Amt proteins was also examined using the strains expressing each Amt protein from their chromosomal loci with the mNeonGreen (mNG) tag at the C-terminus. The Amt1-mNG fusion protein was detected as punctate signals in the cytosol under nitrogen-rich conditions. After shifting the cells to a growth medium without ammonium, Amt1-mNG was translocated to the plasma membrane, mostly at cellular poles (Fig. 2C). On the other hand, Amt2-mNG was detectable at the plasma membrane as patchy structures even under nitrogen-rich conditions. However, after nitrogen starvation, the Amt2-mNG signal at the plasma membrane appeared to increase further (Fig. 2C). We could not detect the Amt3-mNG signal (data not shown), as predicted from the immunoblotting data (Fig. 2A). These observations indicate that the expression and localization of the Amt proteins are differently regulated.

### TORC1 negatively regulates the Amt1 expression

Both immunoblotting and fluorescent microscopy data (Fig. 2A and 2C) showed the induced expression of Amt1 in response to nitrogen starvation. Consistently, real-time quantitative PCR (RT-qPCR) experiments detected an increase in the *amt1^+^* mRNA level in nitrogen-starved cells, implying the transcriptional regulation of *amt1^+^* in response to ammonium (Fig. 3A). Detailed kinetic analysis of the cellular Amt1 protein level found that Amt1 gradually accumulated after nitrogen starvation and that the re-addition of ammonium to the growth medium decreased Amt1 along the time course (Fig. 3B, top panel). In parallel, we assessed the activity of TORC1 by monitoring the phosphorylation of Psk1. As previously reported (Nakashima et al., 2012), ammonium depletion resulted in TORC1 inactivation, which was detected as the loss of the Psk1 phosphorylation (Fig. 3B, middle panel). Upon re-addition of ammonium, the phosphorylation of Psk1 was restored in response to the reactivation of TORC1. Thus, the Amt1 expression appears to negatively correlate with the TORC1 activity.

**Figure 3.**
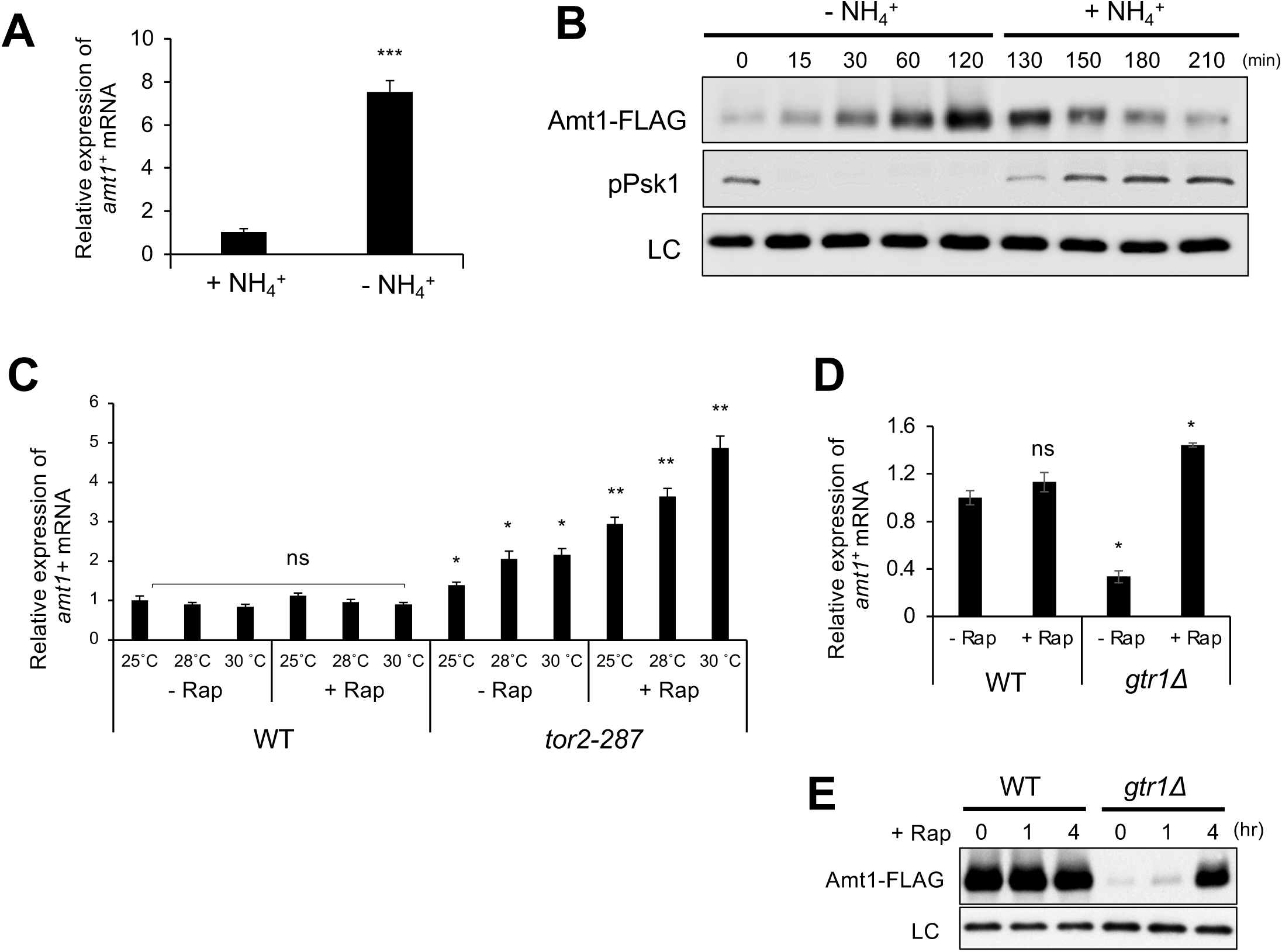
Amt1 expression is negatively regulated by TORC1 signaling pathway. (A) Increased *amt1^+^* mRNA in response to nitrogen starvation. Wild-type cells were cultured in EMM liquid medium at 30℃ and transferred to EMM lacking ammonium. The relative expression of the *amt1^+^*mRNA was normalized to the *act1^+^* mRNA and analyzed in nitrogen-starved cells and nitrogen-rich cells. Mean and SEM of three biological replicates are shown. Asterisks indicate t. test values; p < 0.001 (*\*\*\**). (B) Amt1 protein is negatively regulated by ammonium. Wild-type cells expressing Amt1-FLAG were cultured in EMM at 30 ℃ and transferred to EMM lacking ammonium, followed by re-addition of ammonium to the medium. The crude cell lysates were analyzed by immunoblotting using anti-FLAG, anti-pPsk1 and anti-Spc1 (loading control) antibodies. (C) Inhibition of TORC1 in *tor2-287* mutant increased *amt1^+^* mRNA. Wild-type and *tor2-287* cells exponentially growing at 25 ℃ were shifted to different semi-permissive temperatures (25 ℃, 28 ℃ and 30 ℃), in the absence and presence of rapamycin (200 ng/mL) for one hour. (D) Hyperactivation of TORC1 in *gtr1Δ* mutant decreased *amt1^+^* mRNA. Wild-type and *gtr1Δ* cells exponentially growing in EMM liquid medium at 30 ℃ were treated with rapamycin (200 ng/mL) for one hour. The relative expression of the *amt1^+^*mRNA was normalized to the *act1^+^* mRNA and analyzed in the mutant and wild-type cells, under different treatments. Mean and SEM of three biological replicates are shown. Asterisks indicate t. test values; non-significant (*ns*), p < 0.05 (*\**) and p < 0.01 (*\*\**). (E) Amt1 protein is negatively regulated by TORC1. Wild-type and *gtr1Δ* cells were cultured in EMM at 30 ℃ and transferred to EMM lacking ammonium. The crude cell lysates were analyzed by immunoblotting using anti-FLAG and anti-Spc1 (loading control) antibodies.

To investigate the involvement of TORC1 in the *amt1^+^*regulation, we used the specific TORC1 inhibitor rapamycin. Treatment of wild-type cells with rapamycin did not significantly affect the *amt1^+^*mRNA level (Fig. 3C), possibly due to the incomplete inhibition of *S. pombe* TORC1 by the drug (Takahara and Maeda, 2012). Therefore, we employed the *tor2-287* mutant, which carries a mutation within the catalytic domain of the Tor2 kinase and exhibits rapamycin- and temperature-sensitive TORC1 phenotypes (Hayashi et al., 2007). The *amt1^+^* expression was found to be significantly increased in *tor2-287* cells at the tested temperatures (25°C, 28°C, and 30°C; Fig. 3C). Moreover, treatment of *tor2-287* cells with rapamycin further promoted the expression of the *amt1^+^* mRNA.

In *S. pombe,* the heterodimeric Rag-family GTPases Gtr1-Gtr2 attenuate TORC1 signaling, and the loss of the Gtr heterodimer results in TORC1 hyperactivation (Chia et al., 2017). The *gtr1Δ* mutant displayed decreased expression of the *amt1^+^*mRNA (Fig. 3D) and the Amt1 protein (Fig. 3E), suggesting that hyperactivated TORC1 in this mutant suppresses the *amt1*^+^ expression. Indeed, the cellular levels of both *amt1^+^* mRNA and the protein were restored by attenuating the TORC1 activity with rapamycin (Figs. 3D and E).

Together, these findings strongly suggest that TORC1 negatively regulates the expression of the *amt1*^+^ gene.

### The Gaf1 transcription factor regulates the *amt1^+^* expression

In the budding yeast *S. cerevisiae*, the transcription of the *MEP* and other permease genes is controlled by the GATA-family transcription factors during nitrogen starvation. Upon nitrogen starvation, the Sit4 and PP2A protein phosphatases dephosphorylate the GATA transcription activators Gln3 and Gat1, which translocate into the nucleus and induce the permease genes required to uptake nitrogen-containing nutrients (Georis et al., 2011). The GATA transcription factors Gaf1 and Fep1 in fission yeast have zinc finger motifs and DNA-binding domains similar to those of Gat1 and Gln3 in budding yeast (Fig. S2). To examine whether Gaf1 and/or Fep1 regulate the expression of *amt1^+^*, we quantified the *amt1^+^* mRNA level in the *gaf1Δ* and *fep1Δ* mutant strains by RT-PCR. Under nitrogen starvation, the *amt1^+^* mRNA was induced in both wild-type and *fep1Δ* cells but not in *gaf1Δ* cells (Fig. 4A). Consistently, the nitrogen starvation-induced accumulation of the Amt1 protein was abolished in the *gaf1*Δ strain (Fig. 4B), suggesting that the transcription activator Gaf1 is essential for the induction of *amt1^+^* in response to nitrogen starvation.

**Figure 4.**
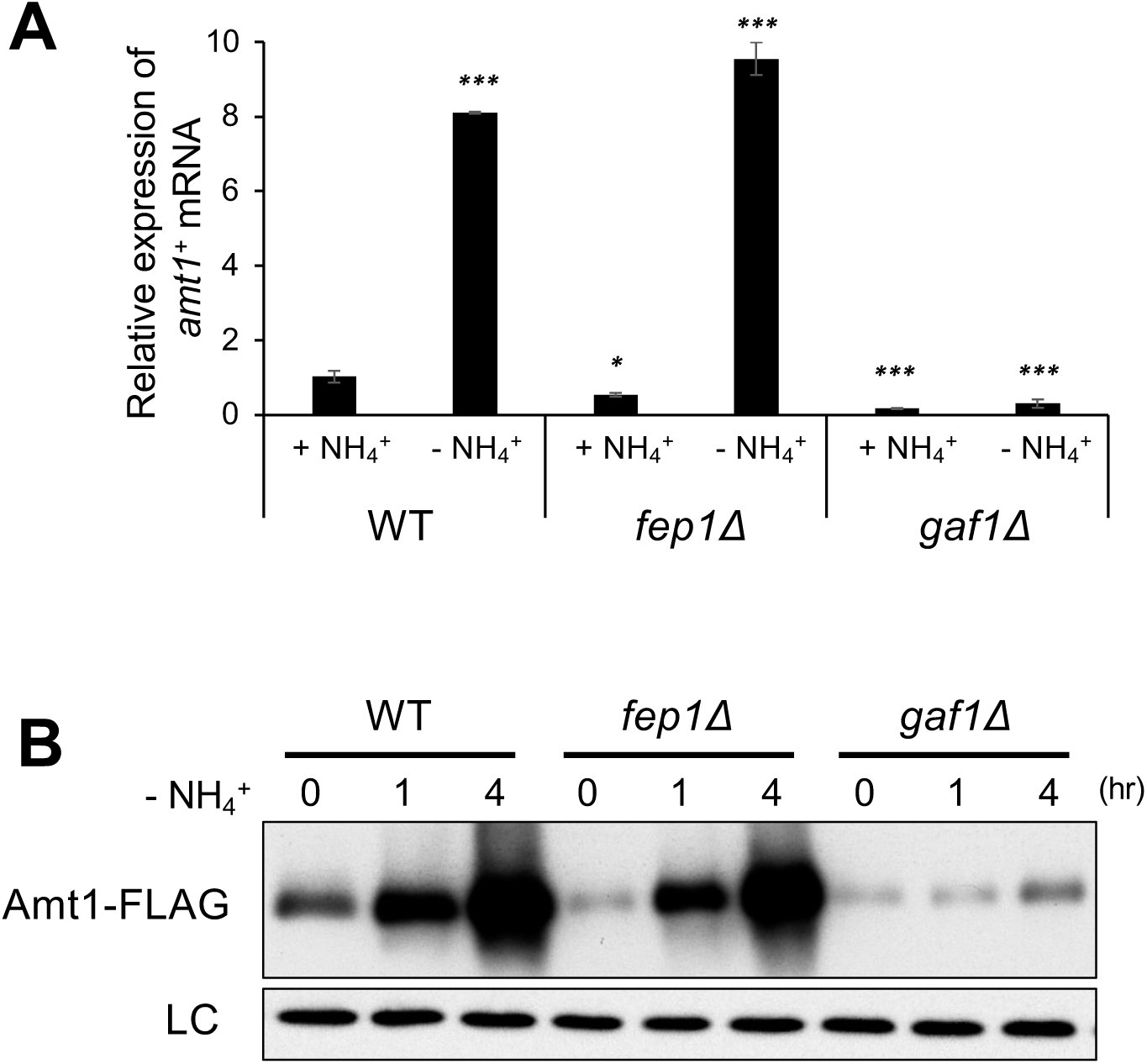
*amt1^+^* mRNA expression is controlled by GATA transcription factor Gaf1 in response to nitrogen starvation. (A) Gaf1 is important for *amt1^+^* induction in response to nitrogen starvation. Wild-type, *fep1Δ* and *gaf1Δ* cells were cultured in EMM liquid medium at 30 ℃ and transferred to EMM lacking ammonium for one hour. The relative expression of the *amt1^+^* mRNA was normalized to the *act1^+^* mRNA and analyzed in the mutant versus wild-type cells; under nitrogen-rich and -starved conditions. Mean and SEM of three biological replicates are shown. Asterisks indicate t. test values; p < 0.05 (*\**) and p < 0.001 (*\*\*\**). (B) Gaf1 is essential for Amt1 accumulation during nitrogen starvation. Wild-type, *fep1Δ* and *gaf1Δ* cells expressing Amt1-FLAG were cultured in EMM liquid medium at 30 ℃ and transferred to EMM lacking ammonium. The crude cell lysates were analyzed by immunoblotting using anti-FLAG and anti-Spc1 (loading control) antibodies.

### Ppe1 phosphatase contributes to the *amt1^+^* expression by Gaf1

It has been reported that Gaf1 is dephosphorylated by the Ppe1 phosphatase and translocates to the nucleus when fission yeast cells are starved of nitrogen (Laor et al., 2015; Ma et al., 2015). Consistently, we observed that Gaf1-GFP translocated into the nucleus upon nitrogen starvation in wild-type but not *ppe1Δ* cells (Fig. 5A). In addition, the TORC1 inhibition by rapamycin induced the nuclear accumulation of Gaf1 even in the presence of nitrogen (Fig. 5B). Moreover, no apparent nuclear accumulation of Gaf1-GFP was detected in *ppe1Δ* cells treated by rapamycin, supporting the model that TORC1 regulates the Gaf1 nuclear localization through Ppe1 (Laor et al., 2015; Ma et al., 2015). It should be noted, however, that rapamycin treatment *per se* does not induce the *amt1*^+^ expression (Fig. 3C) despite the observed nuclear accumulation of Gaf1 (Fig. 5B).

**Figure 5.**
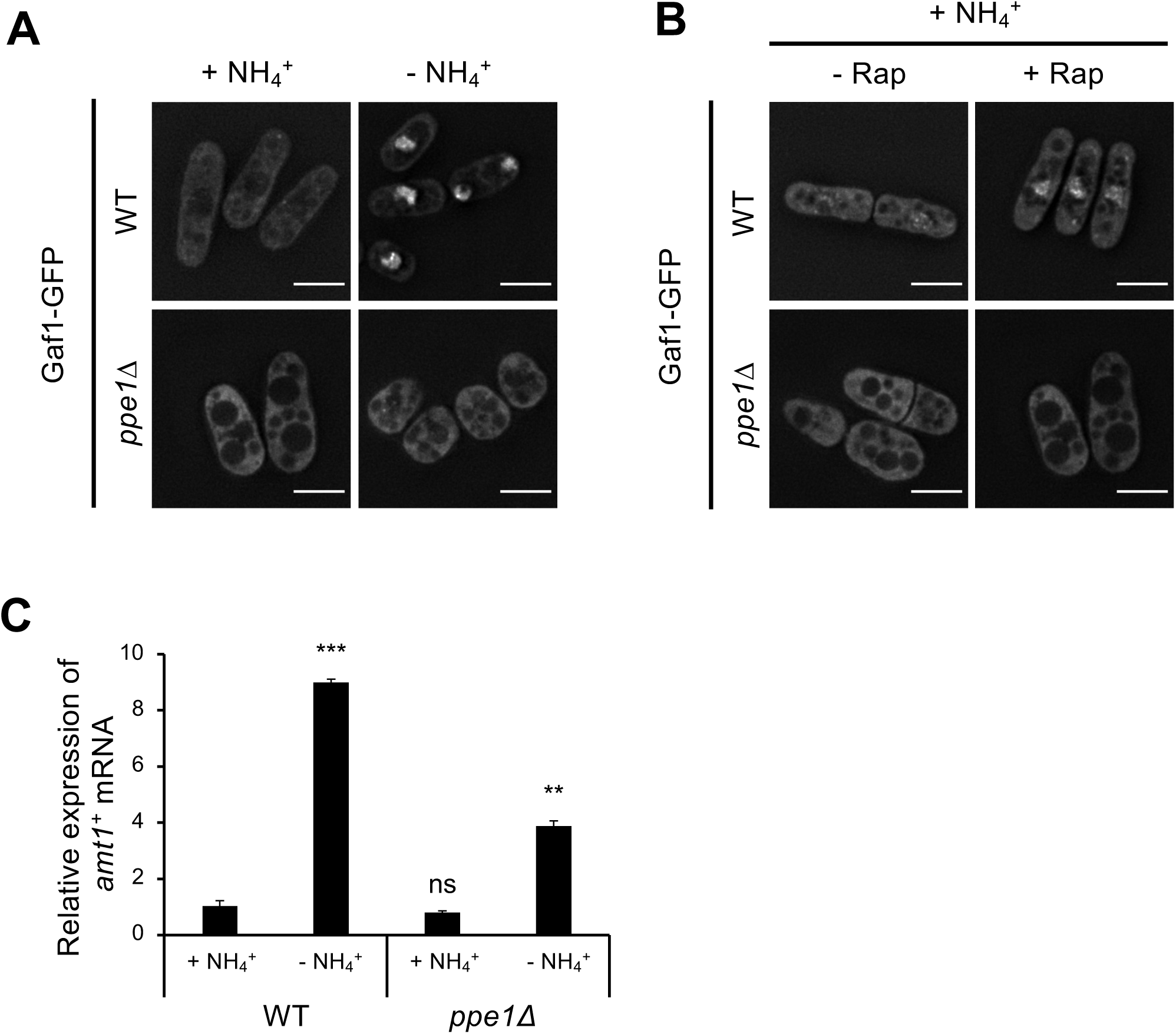
Ppe1 phosphatase regulates *amt1^+^* induction by promoting the nuclear accumulation of Gaf1. Ppe1 is required for Gaf1 nuclear localization during (A) nitrogen starvation and (B) when TORC1 is inhibited by rapamycin. Wild-type and *ppe1Δ* cells expressing the Gaf1-GFP fusion protein were cultured in EMM at 30 ℃. Wild-type and *ppe1Δ* cells were cultured in EMM liquid medium at 30℃ and transferred to EMM lacking ammonium for one hour. scale bar: 5 μm. (C) Ppe1 positively controls *amt1^+^* transcription. Wild-type and *ppe1Δ* cells were cultured in EMM liquid medium at 30℃ and transferred to EMM lacking ammonium for one hour. The relative expression of the *amt1^+^* mRNA was normalized to the *act1^+^* mRNA and analyzed in the mutant versus wild-type cells; under nitrogen-rich and -starved conditions. Mean and SEM of three biological replicates are shown. Asterisks indicate t. test values; non-significant (*ns*) p < 0.01 (*\*\**) and p < 0.001 (*\*\*\**).

Therefore, we next examined whether the Ppe1-dependent nuclear targeting of Gaf1 is required for the *amt1*^+^ induction upon nitrogen starvation. As shown in Fig. 5C, the nitrogen starvation-induced expression of *amt1*^+^ is partially compromised in the *ppe1Δ* mutant compared to that in the wild-type strain. However, a significant increase in the *amt1*^+^ mRNA was still detectable in *ppe1Δ* cells after nitrogen starvation, implying that a Ppe1-independent mechanism controls the *amt1^+^* expression in response to the starvation (Fig.5C).

These results imply that the Ppe1-dependent nuclear accumulation of Gaf1 is not essential for the *amt1*^+^ induction after nitrogen starvation, though such Gaf1 regulation appears to contribute to the induction level partly.

### PP2A is not required for the Gaf1 nuclear accumulation during nitrogen starvation

In fission yeast, the *ppa1^+^* and *ppa2^+^* genes encode the minor and major PP2A catalytic subunits, respectively (Kinoshita et al., 1990). It has been reported that not only *ppe1Δ* but also *ppa1Δ* and *ppa2Δ* affect the phosphorylation state of the Gaf1 protein (Laor et al., 2015). We also consistently observed that the electrophoretic mobility of Gaf1 in the *ppa1Δ* and *ppa2Δ* mutants was slower than that in wild-type cells under nitrogen starvation (Fig. 6A). Thus, the starvation-induced dephosphorylation of Gaf1 is likely to be partially compromised in the absence of the PP2A catalytic subunits.

**Figure 6.**
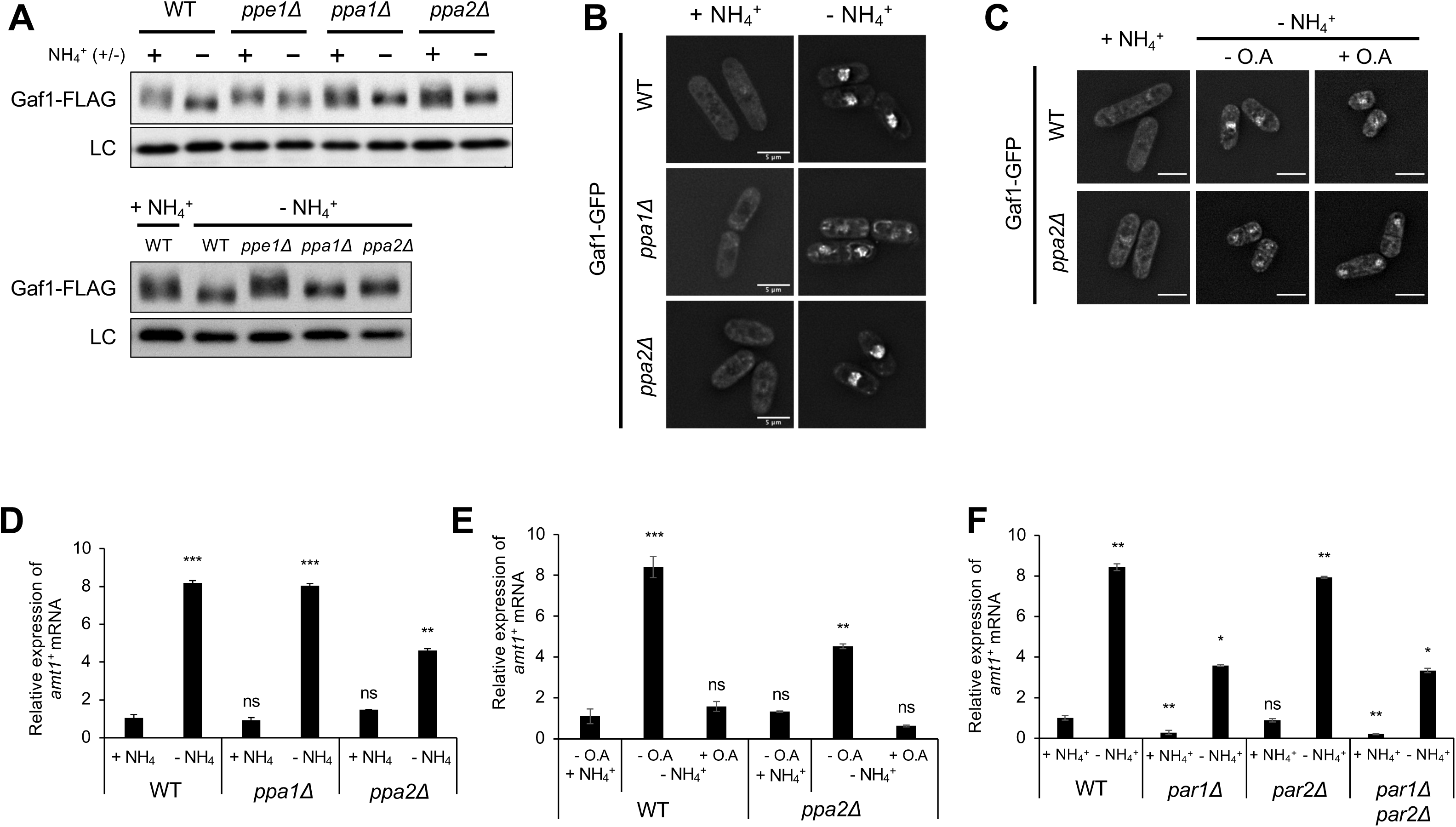
The PP2A-B56^par1^ is essential for the induction of *amt1^+^* through the dephosphorylation of Gaf1. (A-upper) Gaf1 has slower electrophoretic mobility in the deletion mutant compared to wild-type cells during nitrogen starvation. Indicated cells strains expressing Gaf1-FLAG were cultured in EMM liquid medium at 30 ℃ and transferred to EMM lacking ammonium for one hour. The crude cell lysates were analyzed by immunoblotting using anti-FLAG and anti-Spc1 (loading control) antibodies. (A-lower) Side-by-side comparison of the Gaf1 mobility shift in nitrogen-starved cells. (B) PP2A were dispensable for Gaf1 nuclear accumulation. The indicated strains expressing the Gaf1-GFP fusion protein were cultured in EMM at 30 ℃ and transferred to EMM lacking ammonium for one hour. The cells were collected at the indicated time points and observed by a fluorescence microscope. (C) Gaf1 nuclear localization is not affected by PP2A. Wild-type and *ppa2Δ* cells expressing Gaf1-GFP were grown in EMM liquid medium at 30℃ and treated with 20 μM okadaic acid for 30 min. The cells were then washed and transferred to EMM lacking ammonium for one hour with and without 20 μM okadaic acid, and observed by a fluorescence microscope. “-NH_4_^+^” indicates no nitrogen and “+ O.A” indicates addition of okadaic acid. scale bar: 5 μm. (D) Ppa2 positively regulates *amt1^+^* expression. Cells of the indicated strains were cultured in EMM liquid medium at 30℃ and transferred to EMM lacking ammonium for one hour. The relative expression of the *amt1^+^* mRNA was normalized to the *act1^+^* mRNA and analyzed in the mutants versus wild-type cells; under nitrogen-rich and -starved conditions. Mean and SEM of three biological replicates are shown. Asterisks indicate t. test values; non-significant (*ns*), p < 0.01 (*\*\**) and p < 0.001 (*\*\*\**). (E) Okadaic acid treatment abolished the *amt1^+^* induction, in a PP2A-dependent manner. Wild-type and *ppa2Δ* cells cultured in EMM liquid medium at 30 ℃ were treated with okadaic acid (20 μM) for 30 min. The cells were then washed and transferred to EMM lacking ammonium for one hour, which is supplemented with 20 μM okadaic acid. The relative expression of *amt1⁺* mRNA was analyzed as described for Fig. 6D. Asterisks indicate t. test values; non-significant (*ns*), p < 0.01 (*\*\**) and p < 0.001 (*\*\*\**). (F) Regulatory subunit of PP2A, Par 1 regulates *amt1^+^* expression. Cells were treated as described for Fig. 6D. Asterisks indicate t. test values; non-significant (*ns*), p < 0.05 (*\**) and p < 0.01 (*\*\**).

As the Ppe1-dependent dephosphorylation controls the Gaf1 nuclear translocation, we investigated whether the PP2A enzymes are also involved in this event. Under nitrogen-starved conditions, the nuclear accumulation of Gaf1 was observed in *ppa1Δ* and *ppa2Δ* cells as in wild-type cells (Fig. 6B). We hypothesized that this result could be due to the redundant function of Ppa1 and Ppa2. As deleting both *ppa1^+^* and *ppa2^+^* genes results in cell lethality and a viable double deletion mutant cannot be constructed (Kinoshita et al., 1990), we employed okadaic acid, a specific PP2A inhibitor drug (Bialojan and Takai, 1988). A previous study found that 20 µM okadaic acid effectively inhibits the cell division of the *ppa2Δ* mutant, indicating the inhibition of Ppa1 and the complete loss of PP2A activity in the *ppa2Δ* strain under this condition (Kinoshita et al., 1993). We found that 20 µM okadaic acid added to the growth medium did not affect the nuclear accumulation of Gaf1-GFP after nitrogen starvation in both wild-type and *ppa2Δ* cells (Fig. 6C). These observations strongly suggest that, unlike Ppe1, PP2A is dispensable for the Gaf1 nuclear accumulation in response to nitrogen starvation.

### PP2A activity is essential for the *amt1^+^* induction upon nitrogen starvation

Having found that PP2A is not required for the nuclear accumulation of Gaf1 after nitrogen starvation, we next investigated whether PP2A is involved in the starvation-induced expression of *amt1^+^*. RT-qPCR experiments detected comparable induction of the *amt1^+^* mRNA in the *ppa1Δ* and the wild-type cells upon nitrogen starvation, but the induction was significantly reduced in the *ppa2Δ* mutant (Fig. 6D). Thus, the major PP2A catalytic subunit Ppa2 appears to play an important role in the induction of *amt1*^+^ in response to nitrogen starvation.

It is possible that the residual induction of the *amt1^+^* mRNA in *ppa2Δ* cells (Fig. 6D) is mediated by Ppa1. Therefore, we again utilized okadaic acid to inhibit the cellular PP2A activity; in the presence of 20 µM okadaic acid, no apparent induction of *amt1^+^*was detected in both wild-type and *ppa2Δ* strains upon starvation (Fig. 6E) despite the nuclear accumulation of Gaf1 under the condition (Fig. 6C). These findings suggest that the PP2A activity plays an essential role in the transcriptional induction of *amt1^+^* upon nitrogen starvation, independently of the Ppe1-regulated nuclear accumulation of Gaf1.

There are two known regulatory subunits of PP2A, namely the B55 and the B56 subunits (Jiang, 2006). In fission yeast, the *pab1*^+^ gene encodes the B55 subunit, while the paralogous genes *par1*^+^ and *par2*^+^ encode the B56 subunit proteins, with *par1*^+^ carrying the major B56 function (Jiang and Hallberg, 2000). PP2A-B55^pab1^ is active under nutrient-replete conditions (Martin et al., 2017), while PP2A-B56^par1^ is activated during nitrogen starvation (Stonyte et al., 2020). Therefore, we examined whether the B56^par1^ subunit is required for the function of PP2A in the starvation-induced *amt1^+^* expression. In comparison to the wild-type strain, the *amt1^+^* induction upon nitrogen starvation was significantly compromised in the *par1Δ* mutant (Fig. 6F), indicating a significant role of the B56 subunit Par1 in the *amt1^+^* regulation. On the other hand, the *par2Δ* mutation showed little impact on the *amt1*^+^ expression in both *par1*^+^ and *par1Δ* backgrounds.

### The Tsc complex controls the localization of ammonium transporters via Pub1-Any1

In addition to the regulation of TORC1 signaling, the Tsc complex-Rhb1 axis controls the translocation of amino acid transporters to the plasma membrane during nitrogen starvation (Nakase et al., 2013). We discovered that the Tsc complex also controls the localization of the ammonium transporters Amt1 and Amt2 in response to starvation. In wild-type cells, Amt1-mNG translocates from the punctate structures in the cytoplasm to the plasma membrane within 20 minutes after a shift from nitrogen-rich to nitrogen-limited conditions (Fig. 2C). This starvation-induced translocation of Amt1-mNG was significantly compromised in *tsc1Δ* and *tsc2Δ* mutant cells (Fig. 7A). Similarly, the amount of Amt2-mNG at the plasma membrane increased after in wild-type cells starved of nitrogen, but no such increase was observed in the *tsc1Δ* and *tsc2Δ* mutants. These observations suggest that the Tsc1-Tsc2 complex is required for the localization regulation of not only amino acid transporters but also the ammonium transporters Amt1 and Amt2 in response to nitrogen starvation.

**Figure 7.**
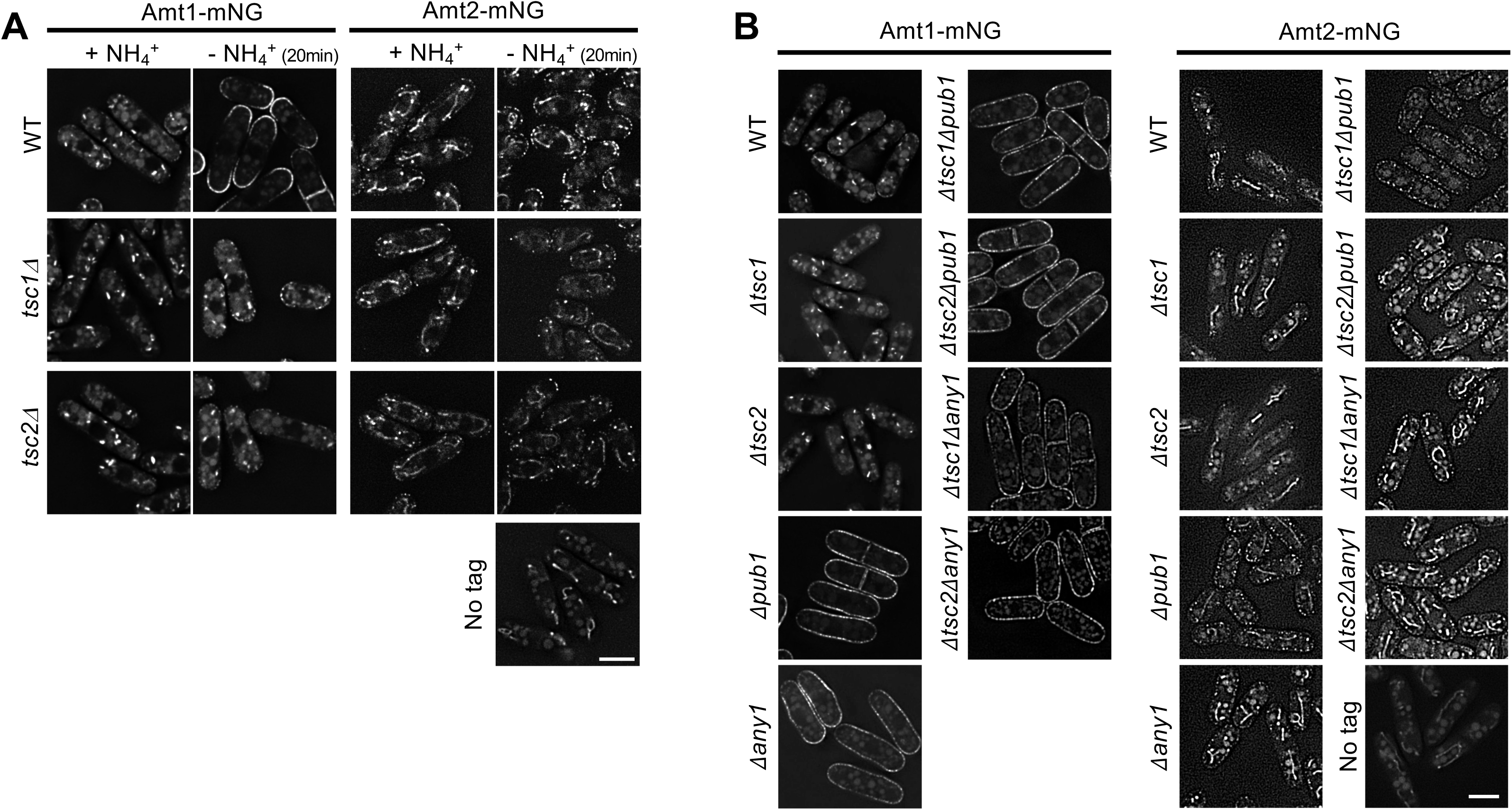
The localization of ammonium transporters is regulated by the Tsc complex through Pub1-Any1. (A) The plasma-membrane localization of Amt1 and Amt2 is compromised in the *tsc1Δ* and *tsc2Δ* mutants. Cells expressing Amt1-mNG or Amt2-mNG were cultured in EMM at 30℃ and transferred to EMM lacking ammonium. The cells were collected 20 min after nitrogen depletion and observed under a fluorescence microscope. Scale bar: 5 μm. (B) Deletion of *pub1* or *any1* restored the plasma-membrane localization of Amt1 and Amt2 that was lost in the *tsc1Δ* and *tsc2Δ* mutants. Cells expressing Amt1-mNG or Amt2-mNG were cultured in EMM at 30℃ and collected at the indicated time points and observed under a fluorescence microscope. Scale bar: 5 μm.

The E3 ubiquitin ligase Pub1 and the β-arrestin-like protein Any1 form a complex that functions downstream of the Tsc-Rhb1 pathway to suppress the plasma-membrane localization of amino-acid transporters in the presence of nitrogen; deletion of either Pub1 or Any1 results in the plasma-membrane localization of amino acid transporters even in the *tscΔ* mutants (Nakase et al., 2013). Observation of Amt1 localization in the *pub1Δ* and *any1Δ* mutants showed that Amt1 localized to the plasma membrane even under nitrogen-rich conditions. Similarly, the amount of Amt2 at the plasma membrane of *pub1Δ* and *any1Δ* mutants was higher than that in the wild type (Fig. 7B). Moreover, the plasma-membrane localization of Amt1 and Amt2 in the *tscΔ pub1Δ* and *tscΔ any1Δ* double mutants was similar to that observed in the *pub1Δ* and *any1Δ* single mutants. Thus, like that of amino-acid transporters (Nakase et al., 2013), the plasma membrane-targeting of the ammonium transporters is likely to be negatively regulated by the Pub1-Any1 complex downstream of the Tsc pathway.

### An unknown ammonium transporter regulated by the Tsc complex

Though the plasma-membrane localization of Amt1 and Amt2 is compromised in the *tsc1Δ* and *tsc2Δ* mutants, their growth was not significantly different from that of the wild type even under the 0.1 mM ammonium condition (Fig. 8A). On the other hand, we found that the *tscΔ* mutations significantly impair the cellular growth under low-ammonium conditions when combined with the *amt1Δ amt2Δ* mutation. Intriguingly, the growth defects of the *amt1Δ amt2Δ tscΔ* triple mutants on 10 mM ammonium medium were much severer than those of the *amt1Δ amt2Δ amt3Δ* triple mutant. Thus, the synthetic phenotype observed between the *amt1Δ amt2Δ* and *tscΔ* mutations is unlikely to be caused by the loss of the Amt3 function.

**Figure 8.**
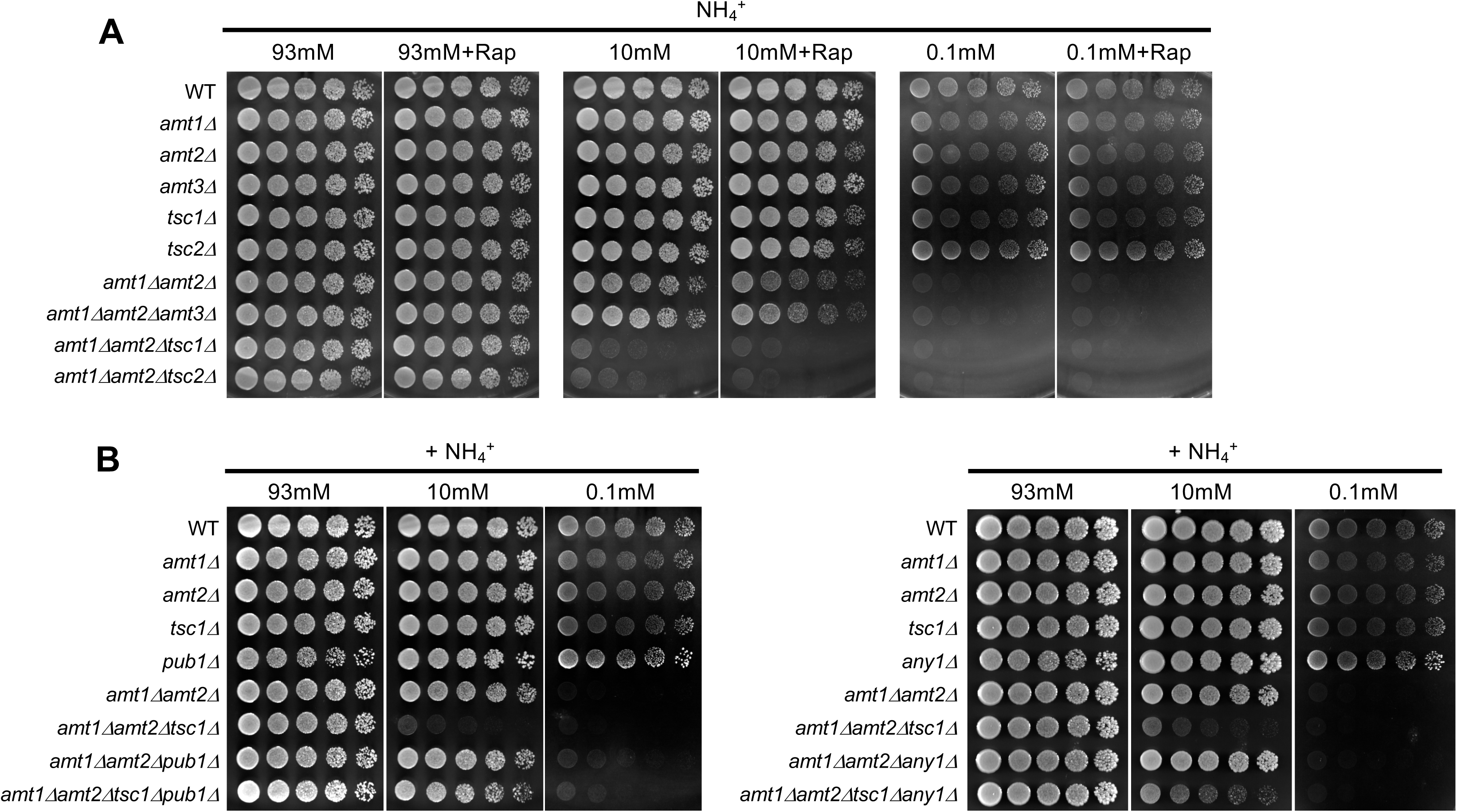
An unidentified ammonium transporter is under the regulation of the Tsc pathway via Pub1-Any1. (A) The combination of *tsc*Δ and *amt1Δ amt2Δ* mutations markedly impairs cellular growth under low-ammonium conditions. Cells of the indicated strains were cultured in EMM medium, and their serial dilutions were spotted onto EMM agar plates supplemented with different concentrations of ammonium (0.1 mM, 10 mM and 93 mM) in the presence or absence of rapamycin, followed by incubation at 30 ℃ for 3 days. (B) Deletion of *pub1* or *any1* suppressed the growth defect of the *amt1Δ amt2Δ tsc1Δ* mutants on 10 mM ammonium medium, but not on 0.1 mM ammonium medium. Cells were treated as described for Fig. 1A.

We confirmed that the hyperactivation of TORC1 in the absence of the Tsc complex (Kwiatkowski, 2003; Manning and Cantley, 2003) is not responsible for the observed *amt1Δ amt2Δ tscΔ* phenotypes, as rapamycin failed to complement the triple mutant growth on 10 mM ammonium, where the *amt1Δ amt2Δ* double mutant can grow (Fig. 8A). Significantly, these growth phenotypes of the *amt1Δ amt2Δ tscΔ* mutants were suppressed by introducing the *pub1Δ* or, to a lesser degree, *any1Δ* mutations; in comparison to the *amt1Δ amt2Δ tscΔ* triple mutants, the *amt1Δ amt2Δ tscΔ pub1Δ* and *amt1Δ amt2Δ tscΔ any1Δ* quadruple mutants grew better on 10 mM ammonium medium (Fig. 8B and Fig. S3). One plausible explanation for these observations is that, like Amt1 and Amt2, another unknown ammonium transporter is under the regulation of the Pub1-Any1 complex and the Tsc pathway (see Discussion).

## Discussion

Through the analysis of the mechanism responsible for the induced expression of the Amt1 ammonium transporter, this study has uncovered previously unknown control of the Gaf1 transcription factor by the PP2A phosphatase. Moreover, our genetic data suggest that, in addition to Amt1, Amt2, and Amt3, fission yeast has yet another ammonium transporter whose plasma membrane localization is under the regulation of the Tsc complex. This study has revealed the intricate modulation of the cellular ammonium uptake in response to the nutrient availability.

Gaf1, a member of the GATA transcription factor family, plays a pivotal role in orchestrating the expression of multiple genes in *S. pombe*. Previous studies have proposed that Gaf1 translocates from the cytoplasm to the nucleus during nitrogen starvation via Ppe1-mediated dephosphorylation, enabling it to both positively and negatively regulate gene expression (Kim et al., 2012; Laor et al., 2015; Rodriguez-Lopez et al., 2020). The *isp7-LacZ* fusion construct in the *ppe1Δ* mutant fails to be induced under a poor-nitrogen condition, suggesting that the Ppe1-dependent nuclear translocation of Gaf1 primarily controls the nitrogen regulation of *isp7*^+^ expression (Laor et al., 2015). While the involvement of PP2A in the dephosphorylation of Gaf1 has been detected, its significance in the regulation of Gaf1 function has never been explored.

In *S. pombe*, Ppa2 serves as the major catalytic subunit of PP2A, and paralogous Ppa1 complements the Ppa2 function (Kinoshita et al., 1993). The lethality of the *ppa1Δ ppa2Δ* double mutant prevents the construction of a complete PP2A knockout, often constraining a comprehensive understanding of the cellular PP2A function. In this study, we have used okadaic acid, a PP2A inhibitor drug, and found that the induced expression of *amt1^+^* upon nitrogen starvation is dependent on the PP2A activity (Fig. 6E). Importantly, under these conditions, the nuclear accumulation of Gaf1 remained unaffected (Fig. 6C). These findings suggest that PP2A along its B56^Par1^ subunit plays a critical role in *amt1*^+^ induction through activation of Gaf1 independently of its nuclear translocation. Thus, Gaf1 appears to be controlled by two distinct phosphatases in response to nitrogen starvation; Ppe1 controls the nuclear targeting, while PP2A-B56^Par1^ governs transcriptional activation. These results have provided a more comprehensive understanding of the spatial and functional regulation of Gaf1-mediated transcription in *S. pombe*.

The regulation of Gaf1 in *S. pombe* shares similarities with the regulation of the GATA transcription factors Gln3 and Gat1 in *S. cerevisiae*. In *S. cerevisiae*, Gln3 and Gat1 activity are controlled by the Sit4 and PP2A phosphatases, both of which operate under TORC1 signaling. Under nitrogen-rich conditions, TORC1 inhibits these phosphatases, while nitrogen starvation relieves this inhibition, leading to the dephosphorylation, nuclear translocation, and activation of Gln3 and Gat1. This demonstrates that TORC1 regulates both localization and transcriptional activity of Gln3 and Gat1 (Di Como and Arndt, 1996; Duvel et al., 2003; Georis et al., 2011; Jiang and Broach, 1999; Tate et al., 2009; Wang et al., 2003; Yan et al., 2006). In *S. pombe*, Ppe1 is known to function downstream of TORC1 (Laor et al., 2015), but whether PP2A-B56^Par1^ is also under TORC1 control remains unclear. However, the marked increase in *amt1**^+^*** expression in rapamycin-treated *tor2-287* cells (Fig. 3C) and the PP2A-dependency of the *amt1*^+^ induction (Fig. 6E) suggest that PP2A also functions downstream of TORC1 in the *amt1*^+^ regulation. Interestingly, recent phosphoproteomic analyses found that Tyr-96 of Par1 undergoes dephosphorylation upon inhibition of TOR signaling (Halova et al., 2021), implying TORC1-mediated regulation of PP2A-B56^Par1^. Identification of the *S. pombe* tyrosine kinase responsible for the Par1 phosphorylation would be of great interest in future studies.

In addition to the transcriptional regulation, this study has also illuminated the localization dynamics of Amt1 in response to nutritional conditions. Under nitrogen-replete conditions, Amt1 localizes intracellularly, while Amt1 exhibits pronounced translocation to the plasma membrane once cells are starved (Figs. 2C & 7). Excessive intracellular ammonium uptake can cause cytotoxicity (Hess et al., 2006; Merrick, 2004), necessitating strict regulation of ammonium transporter activity at the plasma membrane. However, to our knowledge, this is the first report of the regulated localization of ammonium transporters. In *S. cerevisiae,* ammonium transporters Mep1, Mep2, and Mep3 are consistently localized to the plasma membrane, and their activities are regulated by phosphorylation or interaction with inhibitory factors (Boeckstaens et al., 2014; Boeckstaens et al., 2015). In *Escherichia coli*, the activity of the ammonium transporter AmtB is modulated by the binding of the inhibitory PII protein GlnK (Conroy et al., 2007). In *Arabidopsis thaliana*, AMT1;1 is regulated through phosphorylation (Lanquar et al., 2009). It remains possible that any of the ammonium transporters in *S. pombe* are also subject to regulation by phosphorylation or inhibitor proteins.

The regulated localization of Amt1 in *S. pombe* (Fig. 7A) bears striking similarity to that of amino acid transporters, as both are under the control of the Tsc-Rhb1 pathway. The plasma membrane localization of the Aat1 amino acid transporter is regulated downstream of the Tsc-Rhb1 pathway through ubiquitination mediated by the Pub1-Any1 complex (Nakase et al., 2012; Nakase et al., 2013). Intriguingly, recent proteomic analyses (Strachan et al., 2023) detected ubiquitination of Amt1 at Lys-470; considering the involvement of the Pub1-Any1 complex in the control of Amt1 localization (Fig. 7B), it is plausible that, like Aat1, the Amt1 localization is also regulated through ubiquitination by the Pub1-Any1 complex. In contrast, Amt2 exhibited less prominent localization changes under nitrogen starvation. However, significantly altered Amt2 localization in the *tsc* and *pub1* mutant strains (Fig. 7B) implies the Amt2 regulation similar to that of Aat1 and Amt1.

In addition to the confirmed three ammonium transporters Amt1, Amt2, and Amt3, our data has hinted the existence of an unidentified ammonium transporter under the regulation of the Tsc pathway (Fig. 8). As its candidate, we examined *Amf1* (SPBC1683.03c), a *S. pombe* ortholog of the low-affinity ammonium transporter in *S. cerevisiae* (Chiasson et al., 2014). However, its cellular localization and the growth phenotype of the strains lacking Amf1 indicate that Amf1 is not the novel ammonium transporter regulated by the Tsc pathway (Fig. S4). Previous studies in budding yeast proposed that ammonium may enter the cell through K^+^ channels (Hess et al., 2006). There is also evidence that the human aquaporin AQP8 facilitates ammonium transport (Liu et al., 2006). Thus, another Tsc-regulated ammonium transporter to be discovered in fission yeast may have no structural resemblance to the Mep-Amt-Rh family proteins.

Our study has shed light on the multimodal regulation of the *S. pombe* ammonium transporters in response to the nitrogen source. Only Amt1 is subject to TORC1-mediated transcriptional regulation through the Gaf1 transcription factor, which is regulated by the two distinct types of protein phosphatases. Translocation to the plasma membrane is another mode of nutrient-dependent regulation of the ammonium transporters in *S. pombe*. Multiple paralogous ammonium transporters, which were probably acquired by gene duplication, with distinct sets of regulatory mechanisms, are likely to allow cells to fine-tune their response to the nutritional environment. Further studies are expected to delineate such a cellular adaptation strategy developed through evolution.

## Materials and Methods

### Yeast strains and growth media

*S*. *pombe* strains used in this study are listed in Supplementary Table 1. Fission yeast cells were grown at 30 °C in rich medium Yeast extract supplement (YES) and synthetic minimal media, Edinburgh minimal media (EMM) as described previously (Moreno, Klar, and Nurse, 1991; Shiozaki and Russell, 1997). All yeast transformations were performed with the lithium acetate method (Gietz et al., 1992; Okazaki et al., 1990).

### Plasmid construction

The plasmid used in the overexpression studies is a derivative of the pREP1 vector with a thiamine-repressible *nmt1^+^* promoter. Plasmid construction was performed as follows. The pREP1 vector and the *amt1^+^, amt2^+,^* and *amt3^+^* cDNA were cleaved with restriction enzymes *Nde*I and *Not*I, respectively. Following that, both restriction enzyme-digested vectors and cDNA were ligated in which the *amt^+^* was linked downstream of the *nmt1^+^* promoter.

### Gene disruption

Strains were constructed by PCR-based targeting for gene disruption and gene tagging as described before (Bähler et al. 1998). Different types of plasmids harboring different selection markers, respectively, including pFA6A-kanMX6 (kanamycin-resistant), pFA6A-natNT2 (nourseothricin-resistant), and pFA6A-hphMX6 (hygromycin-resistant) were employed for the amplification of PCR-cassettes. The PCR fragments were purified and transformed into fission yeast cells by a simple transformation method as described previously (Morita and Takegawa, 2004). The transformants were selected from selective medium supplemented with G418, nourseothricin, and hygromycin.

### Growth assay

The cells were cultured overnight in EMM/ YES liquid media. The OD_600_ of the culture was measured and adjusted to OD_600_ of 1.0. Serial dilutions of the cells were then spotted on appropriate solid media and allowed to grow for 2-3 days at 30 °C. Images were captured by the Image Quant LAS-4000 (Fujifilm, Japan).

### Measurement of intracellular ammonium concentrations

The cells were cultured to OD_600_ 0.2 in EMM liquid medium containing 10 mM ammonium. Following that, 10 OD cells were collected by filtration (Millipore 0.45 μM pore size filters) and washed 2 times with distilled H_2_O. The filter paper was suspended in a tube containing 1 mL distilled H_2_O, followed by vortexing to dislodge the cells from the filter paper. The cell suspension was boiled at 95 ℃ for 20 min and then subjected to centrifugation at 13,000 rpm for 1 min. Next, 400 µL of the supernatant was transferred to a new tube and stored at -4 ℃, prior to measurement. Concentrations and labeling patterns of intracellular ammonium were subsequently quantified by HPLC as described (Bolten and Wittmann, 2008).

### Protein extraction

The cells were cultured to OD_600_ 0.2 and trichloroacetic acid was added to a concentration of 10 % (v/ v). The cell pellets were harvested by centrifugation at 3,000 rpm for 10 min. Following that, the cell pellets were resuspended with 100 µL 10 % TCA and transferred to a 1.5 mL tube. Zirconia beads were added to meniscus level and subjected to vortexing at 4°C for 10 min. The bottom of the tubes containing cell pellets were then punctured and the cell pellets were collected in a new tube by centrifugation. The cell pellets were resuspended in 150 µL of Laemmli sample buffer and incubated at 65 °C for 10 min. The lysates were then subjected to centrifugation at 13,000 rpm for 5 min and the supernatant was collected in a new tube. The protein concentration of the lysates was measured using Bio-Rad protein assay (Bio-Rad, USA and normalized to 4 µg/ µL before immunoblotting.

### Immunoblotting

Crude cell lysates were prepared as described in the Protein extraction section. Proteins were separated via SDS-PAGE, transferred onto a nitrocellulose membrane, and analyzed using the following primary antibodies: anti-phospho-p70 S6K (1:5000; Cat. No. 9206, Cell Signaling Technology) for detecting anti-phospho-Psk1 (Thr-415) detection, anti-Psk1(1:5000) (Chia et al., 2017), anti-Spc1 (1:10000) (Tatebe and Shiozaki, 2003), and anti-FLAG (1:5000; M2, Sigma-Aldrich). The corresponding secondary antibodies were HRP-conjugated anti-rabbit IgG (H+L) (1:10000; Cat. No.W4011, Promega) and HRP-conjugated anti-mouse IgG (H+L) (1:10000; Cat. No. 112-035-110, Jackson Immuno Research).

### 11-phosphatase treatment

Cell lysates were diluted to 20-30 μg of protein in 1x phosphatase buffer and incubated with 5 mM MnCl_2_ and λ protein phosphatase (λ-PPase) at 30 °C for an hour.

### Fluorescence microscopy

The cells were grown in EMM liquid media to the exponential phase (OD_600_= 0.2). One mL of the cells was collected by centrifugation and resuspended in 100 µL of EMM medium. For the cell preparation on glass slides, agarose was dissolved by boiling in EMM medium to a final concentration of 1.5 % (v/ v). The agarose mixture was prepared by pipetting 20 µL of the mixture on a glass slide and covered with clean plastic film immediately. The cells were mounted on the thin agarose layer before covering it with glass. Fluorescent images were captured using DeltaVision Elite system (GE Healthcare Life Sciences, USA) using x 100 objective lens. Images were deconvolved using the softWoRx software (Applied Precision Inc., USA). The image processing was carried out using ImageJ software.

### RNA extraction and real-time quantitative PCR

The cells were grown in EMM liquid media to exponential phase (OD_600_= 0.2) and collected by centrifugation at 3,000 rpm for 3 min. The supernatant was discarded and the cell pellet was flash frozen by liquid nitrogen. RNA extraction was carried out using the hot phenol method. The cell pellet was mixed with 200 µL bead buffer (75 nM NH_4_OAc and 10 mM EDTA (pH8)) and chloroform isoamyl alcohol. The mixture is mixed with a pre-heated solution comprising 25 µL of 10% SDS, 300 µL acid phenol: chloroform, and acid-washed glass beads that were added to meniscus level. The cell mixture was subjected to a series of 1-min vortex- 1- min heating at 65 °C for 3 times. The samples were then incubated at 65 °C for 10 min and vortexed for 1 minute. Following that, the mixture was centrifuged at 13,000 rpm for 10 min and the aqueous phase was transferred to a pre-heated 250 µL bead buffer (65 °C). The mixture was again centrifuged and the supernatant was transferred to a new tube containing pre-chilled bead buffer with isopropanol. The mixture was centrifuged at 4 °C for 30 min, and the pellets were washed with RNase-free 70 % ethanol. The pellet was centrifuged, dried at room temperature, and finally eluted with 30 µL of RNase-free water.

For genomic DNA removal and cDNA conversion, the PrimeScript RT reagent kit with gDNA eraser (Takara Bio, Japan) was used and carried out according to the manufacturer’s protocol. RT-qPCR was performed using Syber Green MM (Applied Biosystem, USA). The actin1 (*act1^+^*) gene was used as a housekeeping gene for all the RT-qPCR experiments. The nucleotide sequences of the primer sets used in this study are listed in Supplementary Table 2.

## Acknowledgments

We thank Asako Higuchi for her technical assistance and Dr. Hisashi Tatebe for the discussion. We are also grateful to Dr. Tomoyuki Fukuda, Dr. Tomohiro Matsumoto, and the NBRP Japan for the strains.

## Competing Interests

The authors declare no competing interests.

## Author Contributions

Y.N., Y.M., H.T., and K.S. designed research; Y.N., N.S.L., A.S., T.Y., and D.W. performed research; Y.N., N.S.L., and D.W. analyzed data; and Y.N. and K.S. wrote the paper.

## Finding

This study was supported by research grants from the Takeda Science Foundation to K.S. and Y.M., the Tojuro Iijima Foundation for Food Science and Technology to Y.N., and the Institute for Fermentation, Osaka (IFO), and Sumitomo Foundation to Y.M. This work was also supported by the Japan Society for the Promotion of Science (JSPS) KAKENHI grants (20K06485 to Y.N., 19K06564 and 22K06145 to Y.M., 19H03224 to K.S.).

## Notes

### Competing Interest Statement

The authors have declared no competing interest.

