## Supplemental for "Multiple ammonium transporters in fission yeast are coordinated by transcriptional and localization regulation in response to nitrogen starvation": Supplemental information.docx

Running title: Regulation of ammonium transporters by TORC1 in fission yeast

§These authors contributed equally to this work.

*Author for correspondence:

Yukiko Nakase

**Supplementary Table 1: Fission yeast strains used in this study**

| Strain ID | Genotype | Sorce |
| --- | --- | --- |
| CA1 | *h^-^* | Laboratory stock |
| CA101 | *h^-^ leu1* | Laboratory stock |
| CA14576 | *h^-^ amt1∆:: kanMX6* | This study |
| CA14578 | *h^-^ amt2∆:: hph* | This study |
| CA14580 | *h^-^ amt3∆:: nat* | This study |
| CA14582 | *h^-^ amt2∆:: hph amt3∆:: nat* | This study |
| CA14633 | *h^-^ amt1∆:: kanMX6 amt2∆:: hph* | This study |
| CA14635 | *h^-^ amt1∆:: kanMX6 amt3∆:: nat* | This study |
| CA14641 | *h^+^ amt1∆:: kanMX6 amt2∆:: hph amt3∆:: nat* | This study |
| CA11171 | *h^-^ amt1: FLAG (kanMX6)* | This study |
| CA11175 | *h^-^ amt2: FLAG (kanMX6)* | This study |
| CA12271 | *h^-^ amt3: FLAG (kanMX6)* | This study |
| CA11173 | *h^-^ amt1: mNG (kanMX6)* | This study |
| CA11177 | *h^-^ amt2: mNG (kanMX6)* | This study |
| CA11315 | *h^+^ ppe1∆:: kanMX6* | This study |
| CA11119 | *h^-^ gaf1∆:: hph* | This study |
| CA16078 | *h^-^ fep1∆:: kanMX6* | This study |
| CA10985 | *h^-^ gaf1: FLAG (kanMX6)* | This study |
| CA11072 | *h^-^ gaf1: GFP (kanMX6)* | This study |
| CA16086 | *h^-^ amt1: FLAG (kanMX6) fep1∆:: kanMX6* | This study |
| CA16244 | *h^-^ amt1: FLAG (kanMX6) gaf1∆:: hph* | This study |
| CA16305 | *h^-^ gaf1: GFP (kanMX6) ppe1∆:: kanMX6* | This study |
| CA16304 | *h^+^ gaf1: FLAG (kanMX6) ppe1∆:: kanMX6* | This study |
| CA13053 | *h^-^ ppa1∆:: ura4^+^ ura4-D18* | This study |
| CA13055 | *h^-^ ppa2∆:: ura4^+^ ura4-D18* | This study |
| CA17245 | *h^-^ gaf1: FLAG (kanMX6) ppa1∆::ura4^+^ ura4-D18* | This study |
| CA17246 | *h^-^ gaf1: FLAG (kanMX6) ppa2∆:: ura4^+^ ura4-D18* | This study |
| CA17332 | *h gaf1: GFP (kanMX6) ppa2∆:: ura4^+^ ura4-D18* | This study |
| CA18240 | *h gaf1: GFP (kanMX6) ppa1∆:: ura4^+^ ura4-D18* | This study |
| CA8413 | *h^+^ tor2-ts287* | Laboratory stock |
| CA17628 | *h^-^ gtr1∆:: nat* | Laboratory stock |
| CA14829 | *h^-^ amt1: FLAG (kanMX6) gtr1∆:: nat* | This study |
| CA6903 | *h^-^ par1∆:: kanMX6 leu1-32* | This study |
| CA6906 | *h^-^ par2∆:: kanMX6 leu1-32* | This study |
| CA7082 | *h^-^ par1∆:: kanMX6 par2∆:: kanMX6 leu1-32* | This study |
| CA17610 | *h^-^ tsc1∆:: hph* | Laboratory stock |
| CA9055 | *h^-^ tsc2∆:: hph* | Laboratory stock |
| CA10361 | *h^-^ pub1∆:: hph* | Laboratory stock |
| CA11491 | *h^-^ any1∆:: hph* | Laboratory stock |
| CA19802 | *h^-^ amt1: mNG (kanMX6) tsc1∆:: hph* | This study |
| CA12376 | *h^-^ amt1: mNG (kanMX6) tsc2∆:: hph* | This study |
| CA19899 | *h^-^ amt2: mNG (kanMX6) tsc1∆:: hph* | This study |
| CA19901 | *h^-^ amt2: mNG (kanMX6) tsc2∆:: hph* | This study |
| CA17649 | *h^-^ amt1: mNG (kanMX6) pub1∆:: hph* | This study |
| CA19913 | *h^-^ amt1: mNG (kanMX6) any1∆:: hph* | This study |
| CA19907 | *h^-^ amt2: mNG (kanMX6) pub1∆:: hph* | This study |
| CA19909 | *h^-^ amt2: mNG (kanMX6) any1∆:: hph* | This study |
| CA20042 | *h^+^ amt1: mNG (kanMX6) tsc1∆:: hph pub1∆:: hph* | This study |
| CA20050 | *h^-^ amt1: mNG (kanMX6) tsc2∆:: hph pub1∆:: hph* | This study |
| CA20039 | *h^-^ amt1: mNG (kanMX6) tsc1∆:: hph any1∆:: hph* | This study |
| CA20046 | *h^-^ amt1: mNG (kanMX6) tsc2∆:: hph any1∆:: hph* | This study |
| CA20037 | *h^+^ amt2: mNG (kanMX6) tsc1∆:: hph pub1∆:: hph* | This study |
| CA20048 | *h^-^ amt2: mNG (kanMX6) tsc2∆:: hph pub1∆:: hph* | This study |
| CA20041 | *h^-^ amt2: mNG (kanMX6) tsc1∆:: hph any1∆:: hph* | This study |
| CA20044 | *h^+^ amt2: mNG (kanMX6) tsc2∆:: hph any1∆:: hph* | This study |
| CA19799 | *h^+^ amt1∆:: kanMX6 amt2∆:: hph tsc1∆:: hph* | This study |
| CA19508 | *h^-^ amt1∆:: kanMX6 amt2∆:: hph tsc2∆:: hph* | This study |
| CA19868 | *h^-^ amt1∆:: kanMX6 amt2∆:: hph pub1∆:: hph* | This study |
| CA19866 | *h^-^ amt1∆:: kanMX6 amt2∆:: hph tsc1∆:: hph pub1∆:: hph* | This study |
| CA19903 | *h^-^ amt1∆:: kanMX6 amt2∆:: hph any1∆:: hph* | This study |
| CA19905 | *h^-^ amt1∆:: kanMX6 amt2∆:: hph tsc1∆:: hph any1∆:: hph* | This study |
| CA19864 | *h^-^ amt1∆:: kanMX6 amt2∆:: hph tsc2∆:: hph pub1∆:: hph* | This study |
| CA19911 | *h^-^ amt1∆:: kanMX6 amt2∆:: hph tsc2∆:: hph any1∆:: hph* | This study |
| CA20250 | *h^-^ amf1: mNG (kanMX6)* | This study |
| CA20289 | *h^-^ amf1: mNG (kanMX6) tsc1∆:: hph* | This study |
| CA20336 | *h^-^ amf1: mNG (kanMX6) pub1∆:: hph* | This study |
| CA16081 | *h^-^ amf1∆:: kanMX6* | Boineer |
| CA20020 | *h^-^ amt1∆:: kanMX6 amt2∆:: hph amf1∆:: kanMX6* | This study |
| CA20018 | *h^+^ amt1∆:: kanMX6 amt2∆:: hph amt3∆:: nat amf1∆:: kanMX6* | This study |

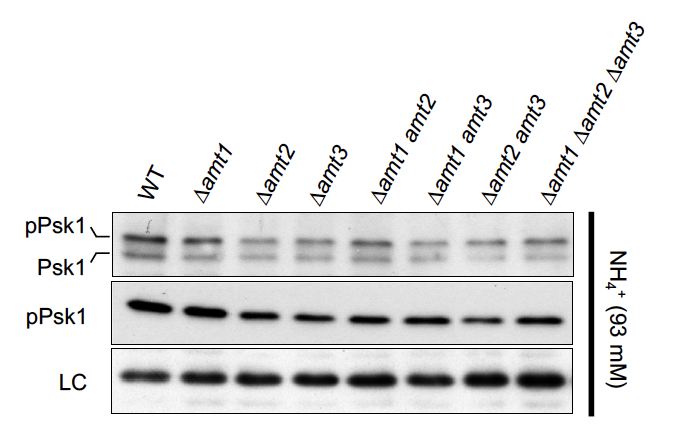

**Supplementary Figure S1. Under 93 mM ammonium, loss of amt genes does not lead to TORC1 inhibition.**

Cells of the indicated strains were grown in EMM medium supplemented with 93 mM ammonium at 30°C. The crude cell lysates were analyzed by immunoblotting using anti-Psk1, anti-pPsk1, and anti-Scp1 (loading control) antibodies.

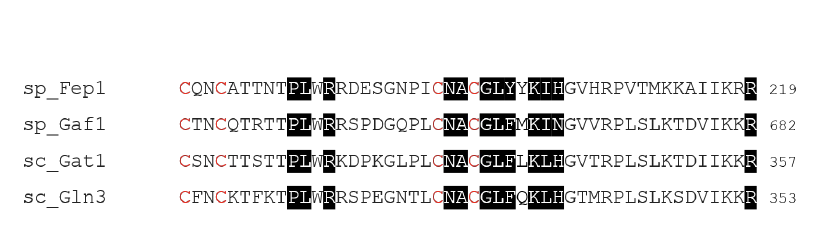

**Supplementary Figure S2. Sequence alignment of the zinc finger motifs in the GATA transcription factors in budding yeast and fission yeast.**

Feb1 and Gaf1 are the GATA transcription factor orthologs in *S. pombe*, and Gat1 and Gln3 are those in *S. cerevisiae*. The four conserved cysteine residues characteristic of the zinc finger motif are indicated in red, and the conserved residues within the DNA-binding domains of the GATA family transcription factors are highlighted in black.

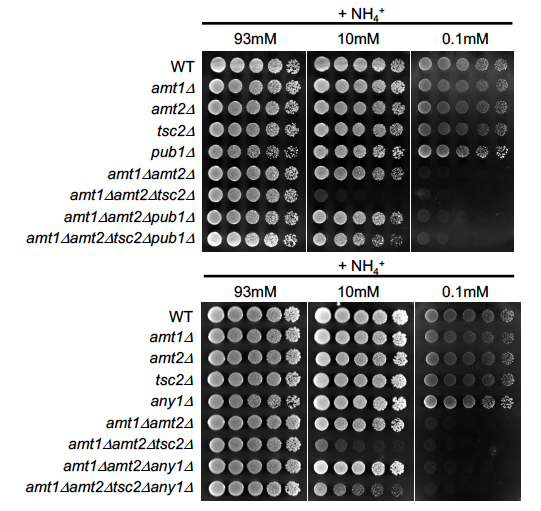

**Supplementary Figure S3. Deletion of pub1 or any1 suppressed the growth defect of the amt1Δ amt2Δ tsc2Δ mutants on 10 mM ammonium medium, but not on 0.1 mM ammonium medium.**

Cells of the indicated strains were cultured in EMM medium, and their serial dilutions were spotted on EMM agar plates supplemented with different concentrations of ammonium (0.1 mM, 10 mM, and 93 mM) and further incubated at 30°C.

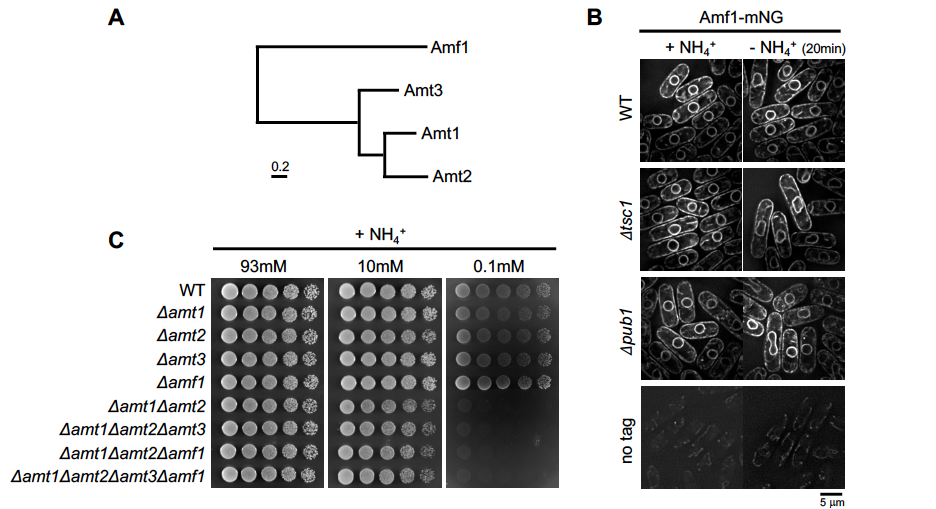

**Supplementary Figure S4. Amf1 is not regulated by the Pub1-Any1 complex downstream of the Tsc pathway.**

(A) Phylogenetic tree of Amt1, Amt2, Amt3, and Amf1 using the ClustalW program. (B) Localization of Amf1 in the presence or absence of nitrogen. Cells expressing Amf1-mEGFP were cultured in EMM at 30 °C and transferred to EMM lacking ammonium for 20 min. (C) Cells of the indicated strains were cultured in EMM medium, and their serial dilutions were spotted on EMM agar medium supplemented with different concentrations of ammonium (0.1 mM, 10 mM, and 93 mM) and further incubated at 30 °C.
